# Teaching R in the undergraduate ecology classroom: approaches, lessons learned, and recommendations

**DOI:** 10.1101/666768

**Authors:** Linda A. Auker, Erika L. Barthelmess

**Affiliations:** Department of Biology, St. Lawrence University, Canton, NY 13617; Department of Biology, Misericordia University, Dallas, PA 18612

**Keywords:** data analysis, data management, pedagogy, R, repeatability, statistics, teaching

## Abstract

Ecology requires training in data management and analysis. In this paper, we present data from the last 10 years demonstrating the increase in the use of R, an open-source programming environment, in ecology and its prevalence as a required skill in job descriptions. Because of its transparent and flexible nature, R is increasingly used for data management and analysis in the field of ecology. Consequently, job postings targeting candidates with a bachelor’s degree and a required knowledge of R have increased over the past ten years. We discuss our experiences teaching undergraduates R in two advanced ecology classes using different approaches. One approach, in a course with a field lab, focused on collecting, cleaning, and preparing data for analysis. The other approach, in a course without a field lab, focused on analyzing existing data sets and applying the results to content discussed in the lecture portion of the course. Our experiences determined that each approach had strengths and weaknesses. We recommend that above all, instructors of ecology and related subjects should be encouraged to include R in their coursework. Furthermore, instructors should be aware of the following: learning R is a separate skill from learning statistics; writing R assignments is a significant time sink for course preparation; and, there is a tradeoff between teaching R and teaching content. Determining how one’s course fits into the curriculum and identifying resources outside of the classroom for students’ continued practice will ensure that R training is successful and will extend beyond a one-semester course.

## Introduction

Thorough training in ecology requires instruction in data management and analysis (Borer et al 2009, Kloser et al 2013, Stevenson et al 2014, Klug et al 2017). For the purposes of this paper, data management skills include data access (via nonproprietary formats and hardware), data organization, and quality control (e.g. looking for data entry errors); analysis skills include exploring data, choosing and applying appropriate statistical tests, converting data to graphical representations, and producing reproducible results (Borer et al 2009, Bravo et al 2016). Often, undergraduate ecology students are exposed to data analysis in their coursework; however, proper data management is not often taught in undergraduate ecology courses (Strasser and Hampton 2012). The primary reason given for not including data management in the curriculum is lack of available course time (Strasser and Hampton 2012). Additional reasons include resistance to changing traditional methods, perhaps due to perceived inappropriateness of teaching data management at the undergraduate level, lack of preparation among the students or the instructor, large class sizes, lack of funding or resources, or the expectation that managing datasets is covered in other courses (Aronova et al 2010, Strasser and Hampton 2012). Often, data are presented as “ready for analysis” and the important steps of data verification and exploratory analysis (e.g., summary statistics and examining distributions) are passed over.

Traditionally, we have each taught Microsoft Excel® for basic data organization, and Excel or other proprietary software for teaching data analysis, as have others (Cass and Ismay 2018). Because many high school students have experience with Excel, it is easier to adopt this software in the lab setting due to the relatively shallow learning curve (Cass and Ismay 2018). However, our own experience and the literature have shown that Excel is insufficient for teaching data management and analysis (Cass and Ismay 2018, Nash 2008). Excel does not make a record of the “point and click” operations performed on the data nor does it offer some of the plot types, such as boxplots, that are helpful for exploratory analysis (Nash 2008). Also, output can remain static even when the value of a datum is changed, leading to incorrect display of statistical results and incorrect plots (Nash 2008). Further, Excel does not handle missing values very well, and can lead to problems with Date formats (Nash 2008). Excel does not allow for clear annotation and commenting, which is critical for open science and reproducibility (Toelch and Ostwald 2018). Finally, some of the statistical output is incorrect (McCollough and Heiser 2008). From a pedagogical standpoint, use of Excel can lead to headaches in grading student work and in determining where students need help, because each student may use a different approach and the lack of code or comments does not separate thought processes from procedural steps.

We have both recently begun to use the R programming environment in our teaching. R is an open-source scripted programming language and environment used for statistical and graphical analysis of datasets, with additional packages readily available for extending its functionality (R Core Team 2018). Like other scripted programming languages, R allows researchers to keep a record of data management and analysis, which allows for reproducibility as one can return to the script at a later date and re-execute it (Borer et al 2009). R is free and flexible, and is increasingly used in the field of ecology. Due to the rise in the use of more complicated statistics in ecology (e.g. mixed effects models and Bayesian statistics), use of R has increased as well (Touchon and McCoy 2016). We have been using R and R Studio (RStudio Team 2016), an open-source and free integrated development environment (IDE) for use with R, for our own scholarly data analyses (henceforth, we use “R” to mean R within the R Studio IDE). The rise in the use of R in ecology, coupled with increasing access to large and complex data sets (Strasser and Hampton 2012) and a push for open science and reproducibility in research (Hampton et al 2015, Klug et al 2017) means that we should be exposing undergraduate students to these modern approaches. Though there is potential framework in place to teach R, there are still few publications that directly address the training of undergraduate ecology students in R.

In this paper, we first argue that data analysis using R is important in ecology by quantifying its increased use for statistical analysis in recent journal papers, as well as in job advertisements that require R as a qualification. We then examine our two different approaches to teaching students how to use R, in conjunction with RStudio, to analyze ecological datasets. We identify advantages common to both approaches, strengths and weaknesses of each approach, and lessons learned from our approaches. Finally, we make recommendations for those wishing to integrate R into their ecology course design. From our approach, we endeavor to begin an important conversation and to lay the framework for improving pedagogy in teaching R to ecology undergraduates.

## Methods

### Putting R in Context

To determine the degree to which R is used in the analysis of ecological data, we examined the first two issues of Ecology published in 2008, 2013 and 2018. For each paper that included data analysis, we recorded the statistical software used. For papers in which R was used, we also noted any packages that were identified. To measure the degree to which skills in R programming are expected for post-undergraduate training and employment, we searched the term “R programming” in the archives of the Ecolog-L Listserv - an established resource for available jobs in the ecological community since 1992 (Inouye 2018) - for the periods Dec 1 - May 31 of the years 2007-2008, 2012-2013, and 2017-2018.

### Approaches to Teaching R

In fall 2017, ELB offered Forest Ecology (FE) and in spring 2018, LAA offered Community Ecology (CE). These were the first offerings for both courses at our primarily undergraduate institution. The courses had several features in common (Table 1). Each explicitly indicated development of R skills as a course component in the syllabus. In CE, the goal was to analyze and understand community ecology datasets with R. In FE, the goal was to introduce data management and statistical testing with R in the treatment of datasets collected by students in the field. In both courses, students were assigned the book *Getting Started with R: An Introduction for Biologists* (henceforth “GSWR”; Beckerman et al. 2017) as one of the required texts. Both courses were offered for advanced undergraduates, all of whom had completed our two-semester sequence of introductory biology, most of whom had taken an introductory ecology course, and some of whom had completed an introductory statistics course. Finally, both courses were small (CE N = 8 students, FE N = 11 students).

**Table 1.** Similarities and differences between two upper-level ecology courses each focused on teaching R as a key component of the course.

| Course characteristic | Forest Ecology | Community Ecology |
| --- | --- | --- |
| Introductory Biology sequence (Biology 101 and 102) required | Yes | Yes |
| Biology 221 (Ecology) | Required | Recommended |
| Number of students | 11 | 8 |
| Lab | Yes | No |
| Number of 1° articles read/discussed | 8 | 5-6 (for one assignment there were two groups of students who each read a different but related paper) |
| Presumed statistical knowledge for students entering course | Low; no assumptions made about prior statistical knowledge | Low; Assumed students had a basic understanding of t tests and ANOVA |
| Number of R assignments | 22 “low stakes” and 2 larger assignments | 8 weekly assignments and one semester-long analysis project |
| How were assignments submitted? | Via email as R scripts and Word documents with embedded figures, as applicable to each assignment | Via Assignments tool on Sakai (course management software). I required R script file, completed handout, and graphs (if applicable to the assignment). |

There were important differences between the courses, as well (Table 1). Community Ecology was organized as a twice-weekly seminar course (with no lab) that met for 1.5 hours each class period, whereas FE met for one 1.5 hour class and one 5.5 hour extended lab period each week. Approximately two-thirds of CE students had previous, though limited, experience with R, whereas no students in FE had used R before. In CE, class time was divided between lecture on community ecology concepts and theories, with some time dedicated to in-class activities and discussion as well as assessments (weekly quizzes and one mid-term exam). In FE, the 1.5 hour lecture periods were primarily used for lecture on forest ecology and for lessons in R. We used the 5.5 hour lab sessions, which for the first 2.5 months of the semester were set primarily in local forests, for gathering data on forest structure as well as discussing papers from the primary literature. During the remainder of the semester, the lab period was used both for lecture and for introducing and getting started on the two major data analysis problem sets assigned during the semester.

Because of the distinct structures of our two courses, our approaches to teaching data management and analysis with R were also different, though complementary. In CE, students were given one assignment per week (for a total of eight assignments) ranging from 5-15 points based on a specific task that was tied into the content of the lecture (example assignments from both CE and FE are in Appendix S1). The task for these assignments required each student to use RStudio to import and analyze a data set and answer questions, applying ecological knowledge. For example, when we covered competition and niches in lecture, the R assignment for that week focused on how to use the *spaa* package (Zhang 2016) to calculate niche overlap among MacArthur’s warblers (MacArthur 1958). As the students gained more experience with R, assignments were designed to encourage them to recall how to do steps that they had done before (e.g., import data from a .csv file) rather than explicitly instruct them each time. Therefore, each student was expected to build on previous knowledge as they progressed through the assignments.

Students used the desktop version of R Studio for their analyses after downloading it independently to their computers. All assignments were written by the instructor or adapted from multiple sources, including exercises from textbooks such as Gardener (2014). Students were also assigned readings from GSWR and applied what they learned from those readings to write their own R code for analysis of data relevant to our course. Students were expected to use R for a final project in which they analyzed a dataset in three different ways: first, through appropriate statistical analysis; second, through some form of visual analysis (either a graph or a map); and third, through analysis of community structure (e.g., diversity or niche overlap). The instructor assigned real datasets from the Ecological DataWiki (https://ecologicaldata.org/home) so students could practice data management skills, such as selecting and formatting the data they needed to do their analyses. Each student was required to meet with the instructor twice during the semester; the first time to discuss the three analyses that they planned to do and the second time to show their progress and troubleshoot, if necessary. The final product for this project was a poster that showed the results of their analyses as well as the R code they used to conduct their analyses. The students presented their posters in a symposium format at the end of the semester to an audience of their peers and department faculty.

In FE, R and R Studio were presented early in the semester. During the first R lesson, to motivate further R learning, students imported and worked with their own data, collected from a local forest during the first lab period. Thereafter, we spent less class time devoted to R until the last third of the semester, but each week, students completed 1-4 short “low stakes” R assignments, each worth two points. We worked through most of the GSWR book (Chapters 1 - 6 and 8, of 9 chapters). In each week’s set of assignments, the first was to read the assigned chapter of GSWR and submit an R script showing that the student had worked through the material in the chapter. The later assignments during the same week asked students to use subsets of the data they collected in the field to complete tasks similar to those covered in that week’s GSWR chapter. All homework assignments were submitted as R scripts via email to the instructor. As in CE, toward the start of the semester, assignment instructions were more detailed including, for example, lines of R code that students should use as well as hints. As the semester progressed, assignments became less detailed so that students had to build on prior knowledge in order to complete the assignments. At both mid-semester and at the end of the semester, students completed a problem set based on analyzing aspects of their forest data. Each problem set listed specific required end products (figures, analyses, etc.) that students were asked to produce without any instruction, thus pushing students to gain more independence in managing data, hypothesis testing, data visualization and writing R code. During the last third of the semester, we devoted class or lab time to discuss application of statistical and ecological analyses (e.g. community ordination) with R. The analysis workflow presented over the course of the semester followed a similar focus to GSWR in how to approach a data set and statistical testing, introduced in Chapter 4 (Table 2). In this workflow, the first three steps could be considered “data management” rather than exploratory or statistical data analysis.

**Table 2.** Steps to data analysis workflow presented in Getting Started With R (Beckerman et al. 2017) and adopted as practice in Forest Ecology.

| Step | Description |
| --- | --- |
| 1 | Import data |
| 2 | Summarize and plot data to look for data entry errors, patterns in data, and outliers |
| 3 | Repair any data errors |
| 4 | Plot relationships and formulate expected outcomes of statisti |
| 5 | Run statistical tests and check for assumptions |
| 6 | Interpret results |
| 7 | Generate final figures/tables that are informed by results. |

Most of the low stakes assignments emphasized use of the *dplyr* (Wickham et al. 2018) and *ggplot2* (Wickham 2016) packages for manipulating and plotting data. Assignment instructions emphasized “cleaning” the data, e.g. looking for and correcting mistakes in data entry, dealing with NAs, examining data for outliers, etc. In both of the problem set assignments, students were asked to demonstrate, using their R code, that they had completed all 7 of the GSWR steps in working with their data. This requirement reinforced the need for data management, including cleaning and repair, prior to analysis, steps often left out of the undergraduate curriculum.

## Results

### Putting R in Context

We examined 56, 54 and 44 Ecology papers (154 total) from 2008, 2013 and 2018, respectively. In 2008, only 59% of published papers indicated the software used for data analysis, and only 12% of those papers (N = 4) indicated using R. Both indication of software used for analysis and use of R for published data analyses increased dramatically over time, reaching 92% and 80%, respectively, by 2018 (Figure 1). Authors were also more detailed about how they used R over time. Of the 4 papers that used R in 2008, two specified the packages used, whereas in 2013, of the 34 papers that used R, 21 specified the packages used, and by 2018, of the 29 papers that used R, 27 specified the packages used. Across years, the most commonly used R packages were lme4 (N = 11), vegan (N = 9), and nlme (N = 7). A list of all identified packages and their frequency of use is in Table 3.

**Table 3.** R packages identified as used for analysis in the first two issues of Ecology from 2008, 2013 and 2018 combined. R packages were mentioned 50 times; Number indicates the number of times a package was identified out of that total.

| Package | Number |
| --- | --- |
| lme4 | 11 |
| vegan | 9 |
| nlme | 7 |
| bbmle, lsmeans, picante | 3 |
| ape, ggplot2, home-made packages, MASS, multcomp, piecewiseSEM, unmarked | 2 |
| ade4, adehabitatHR, AICcmodavg, ape, ca, car, casper, changepoint, deSolve, dplyr, drc, geiger, gridExtra, lmPerm, mclust, metafor, MuMin, packfor, phytools, popbio, psych, quantreg, randomForest, raster, Rmark, several, sncf, sp, spacemakeR, stats, stoichcalc, tidyr, unmarked | 1 |

**Figure 1.**
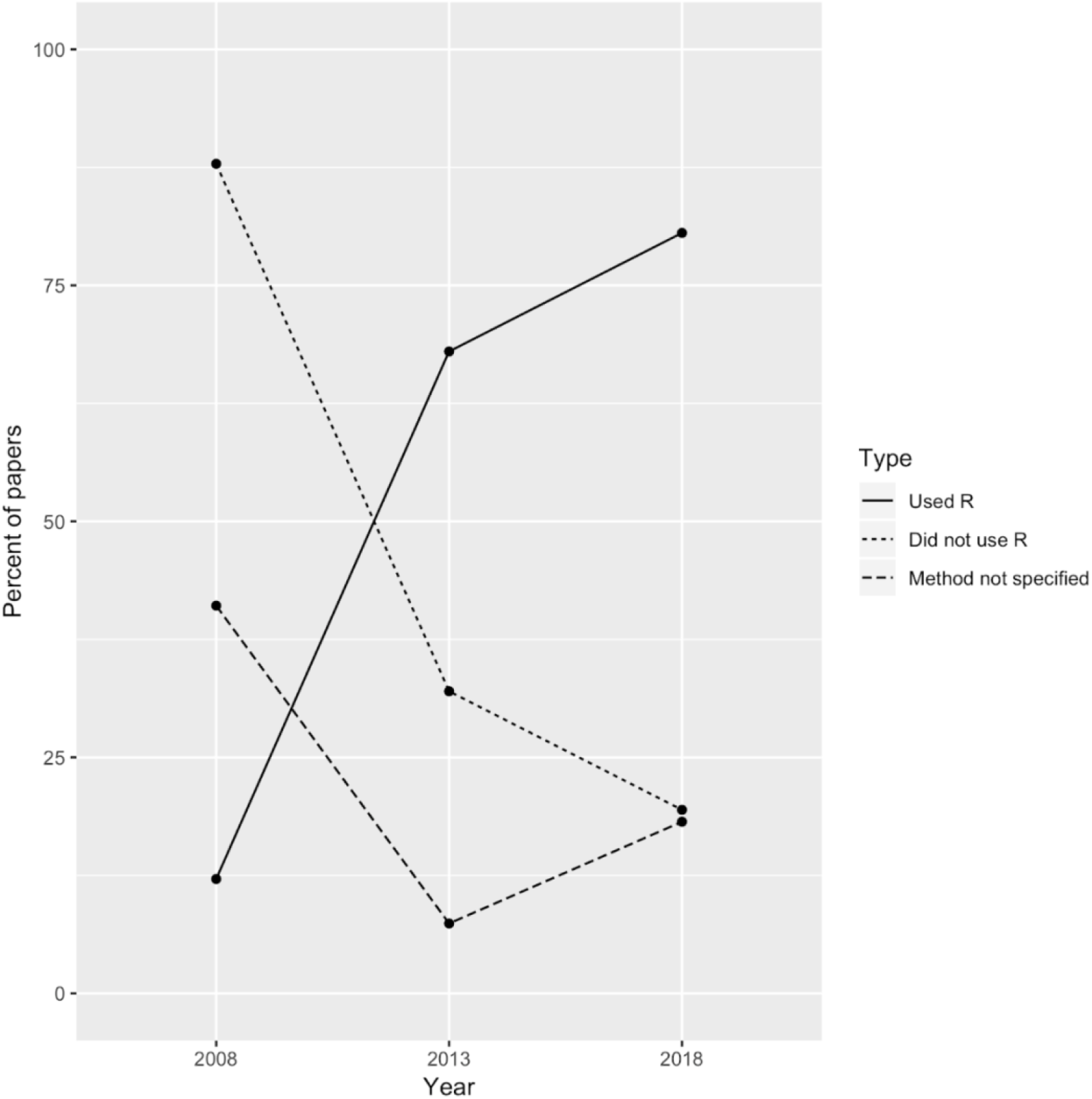
Percent of papers published in the first two issues of Ecology in 2008, 2013 and 2018 in which R was used for data analysis, in which R was not used, and in which the analysis software was not specified.

In terms of the need for R skills for employment, we found that in the 2007-2008 job season, jobs posted on the Ecolog-L listserv requiring R yielded six unique results of which four were postings for postdoctoral opportunities and one was a post for a Ph.D. assistantship. Five years later, in 2012-2013, there were 24 matches for this query: 13 were for postdoctoral opportunities, seven were Ph.D. assistantships, one was a M.S. assistantship, one was a job requiring a postgraduate degree (but not listed as a postdoc), and one was a postgraduate fellowship requiring computation skills, including R. There were no postings for jobs for undergraduates or those who had obtained a bachelor’s degree during this period. Finally, in 2017-2018, there were 47 search results for “R programming”. Of these, four were for a job opportunity (e.g. research technician) that required a bachelor’s degree. There were also 11 posts for Ph.D. assistantships, three posts for M.S. assistantships, and one post for a graduate assistantship with no degree specified. There were 12 posts seeking postdoctoral research associates. There were also two posts for workshops that required previous knowledge of R (one was for distance sampling and not specific to learning more about R) and one post for an internship that both required a bachelor’s degree and knowledge of R. Figure 2 summarizes these findings and shows an increasing requirement for undergraduates in ecology to be able to use R.

**Figure 2.**
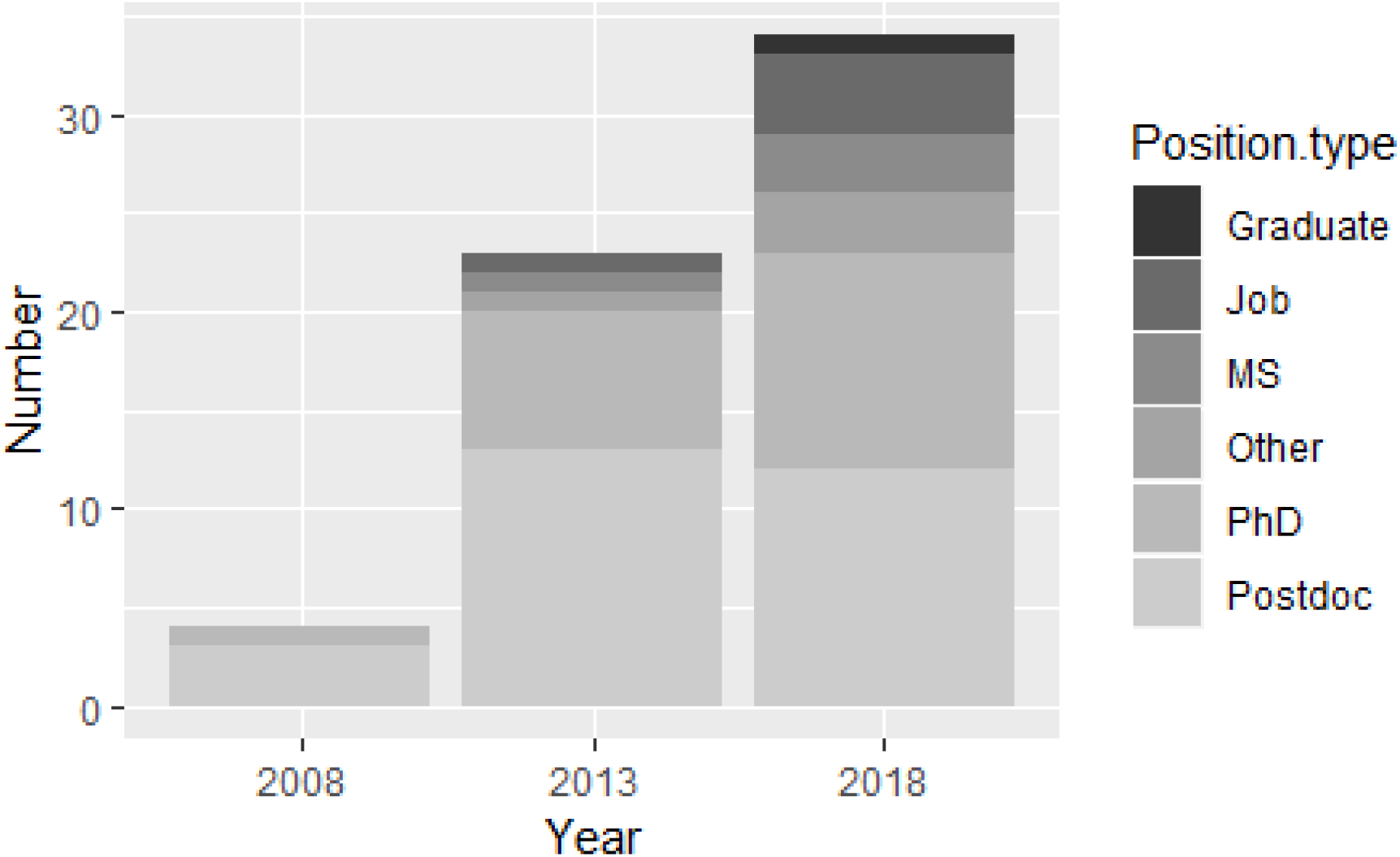
The distribution of position postings on Ecolog-L requiring R for a six-month period Dec 1 – May 31 in 2008-2009, 2012-2013, and 2017-2018. “Graduate” refers to assistantship positions that are not specified as masters or doctoral positions, “Job” refers to a non-degree-seeking position, “MS” is a Masters Program, “Other” is anything requiring R experience not included in the other categories (e.g. workshops), “PhD” is a doctoral program, and “Postdoc” is a postdoctoral program.

### Approaches to teaching R

Students in FE completed 22 low-stakes R assignments and two larger problem sets in which they had to apply their data management and R skills. Students in CE completed 8 weekly R assignments and a large final project. By the end of the semester, students in both courses were confident in their ability to independently import .csv files into R, install and load packages, create and save R scripts, and create and save figures. Students in FE were able to clean and repair datasets and look for outliers prior to analysis. In both classes, some of the students were able to run a series of statistical tests independently, and others with assistance. The list of R skills students developed and R packages students were exposed to are in Table 4.

**Table 4:** R Skills and packages covered in courses (CE = Community Ecology, FE = Forest Ecology, y = used in the course; n = not used). Numbers indicate the number of assignments in which a particular package was used.

|  | Course |  |
| --- | --- | --- |
| R Skill | CE | FE |
| ANOVA | y | y |
| Calculating Niche Overlap | y | n |
| Chi-square | y | y |
| Correlation | n | y |
| Create and save R scripts | y | y |
| Diversity indices | y | y |
| Dissimilarity indices | y | n |
| Dendrograms | y | n |
| Import data | y | y |
| Install and load packages | y | y |
| Installation of R and RStudio | y | y |
| Lotka-Volterra Models | y | n |
| Mapping | y | n |
| Ordination | n | y |
| Plotting with ggplot2 | y | y |
| Regression | y | y |
| Subsetting/Filtering Data | y | y |
| Two sample t test | n | y |
| R package | CE | FE |
| BiodiversityR | 1 | 0 |
| corrgram | 0 | 1 |
| corrplot | 0 | 1 |
| deSolve | 1 | 0 |
| dplyr | 4 | 23 |
| EcoSimR | 1 | 0 |
| ellipse | 0 | 1 |
| ggcorrplot | 0 | 2 |
| ggfortify | 0 | 3 |
| ggmap | 1 | 0 |
| ggplot2 | 3 | 20 |
| ggvegan | 0 | 1 |
| gridExtra | 0 | 1 |
| Hmisc | 0 | 1 |
| raster | 1 | 0 |
| readr | 0 | 24 |
| spaa | 2 | 0 |
| tidyr | 1 | 4 |
| vegan | 2 | 2 |

## Discussion

Our work has shown that R has become the standard tool for ecological data analysis. Further, experience working with R has become a commonly required skill for post-baccalaureate employment and admission to graduate school. However, R has a fairly steep learning curve as a scripted programming language; students with no background in programming may find it more difficult to learn than they would a graphical-user-interface driven software application. Thus, it is imperative that undergraduate programs in biology and ecology begin teaching R to adequately prepare students for the next stages in their careers.

We have presented two different approaches to teaching R in the context of undergraduate ecology courses. In FE, the primary emphasis was on collecting and managing data, with limited statistical analysis, whereas in CE, the primary emphasis was on using R to answer specific community ecology questions with already existing datasets. We found distinct advantages to teaching R regardless of approach and found that there were distinct strengths and weaknesses in each of these two approaches.

Overall, we found that the scientific thought process was reinforced as students made observations from the dataset they were assigned (CE) or which they created (FE) and asked questions they could answer in a stepwise fashion that was clearly traceable in their code. In both courses, students were required to prepare their data for analysis by first fixing mistakes in the data set, removing missing data points (NAs), finding outliers, and subsetting data sets to obtain variables relevant to their question. With R, data management steps such as these can be accomplished in a few lines of code, making them easy to include in teaching. R is far more flexible than a spreadsheet in allowing students to conduct exploratory data analyses, to quickly visualize data, and to look for outliers prior to statistical testing. Further, students can use these visuals to actively predict how a statistical test might turn out, a good practice in scientific thinking. By writing and commenting code, students are able to separate their scientific thoughts from analytical steps, resulting in more clarity of thought regarding their analysis. The process of writing and commenting code as part of an analysis also helps students learn practices in reproducible research. Further, the use of comments and code by students simplified grading of data analysis assignments and aided in troubleshooting problem areas.

### Community ecology strengths and weaknesses

The community ecology (CE) course did not have a laboratory component. As a result, students used published datasets that were freely available. By using these datasets in R, students were given an opportunity to apply their understanding of theory and concepts they learned in lecture, and to see expected patterns within the data. Based on student feedback, the most valuable datasets in terms of student interest and learning were those taken from papers students discussed in the course. In CE, the focus was on the application of R to specific community ecology problems, and so students used specialized packages not covered in the GSWR book.

As far as student attitude and interest in having R introduced in this course, LAA found that by explicitly stating that R would be included in the course description, students who were enrolled expected that learning R would be part of the coursework. In addition, students were encouraged to learn from their errors. For example, in the first lab assignment LAA included an error log to give students a place to record common errors in their code that they could return to later for reference. In-class student reviews about the material were generally positive about learning R, and at least three of the students in the course used R in their senior-year capstone projects.

The consequence of having a course without a lab is that students were limited to already existing data sets. While this saved time so that more theoretical content was covered, students did not have the experience of collecting their own data or practicing good data management for each dataset. However for their final project, students were required to “clean up” a dataset and subset variables, with minimal experience, before they moved on to data analysis. Another result of focusing more on content was that LAA did not spend a lot of time in class going over R assignments. Feedback was mostly limited to comments on assignments submitted via the learning management system, or in one-on-one meetings during office hours. Finally, there are few textbooks that incorporate R into theory in the field. Our textbook was purely conceptual and most of the assignments were adapted by LAA. Therefore, there was a disconnect between readings and hands-on assignments that may be better integrated with a text that uses R to work through relevant community ecology problems.

### Forest ecology strengths and weaknesses

As with CE, students knew from the outset of Forest Ecology that learning R would be a focus of the course. Students used R to manage and analyze data they had collected themselves. A benefit of this course design was that students had the opportunity to directly relate their field observations to the data management and analysis process. When they saw, for example, that the factor variable of “Tree Species” included 4 different versions of “sugar maple,” they were able to easily understand that the error was the result their own errors in data entry and not an abstract problem. Further, because of their connection to the forests, there was a strong motivation for learning R for data analysis to better understand the patterns and processes the students had been observing in the field. This motivation was particularly helpful when the analysis being performed introduced a new concept. For example, near the end of the semester we compared the forests via ordination with the ‘vegan’ package. Ordination is a multivariate technique that is generally not included in introductory statistics classes. Reducing multivariate data sets was thus not familiar to these students. By the end of the semester, however, their familiarity with R allowed us to focus less on the technical side of how to do the ordination, and more on the conceptual side of how to understand what the results of the ordination meant. Familiarity with the forests from which the data were collected allowed the students to consider the results of the ordination relative to their personal experience with each forest, adding an element of understanding.

The primary weakness of this course design was the loss of time devoted to conceptual content. The extended field time meant less lecture time; dividing lecture time between content and learning R meant that we covered less forest ecology content in less depth. Student reviews were positive, both in terms of learning R and in terms of learning field skills; some students observed a desire to have learned more course content. That five of the 11 students in FE opted to enroll in a course to expand their R skills the following semester is testament to the fact that students found value in their growing ability in R.

### Recommendations to other ecology instructors

Because R is becoming increasingly more prevalent in the ecology field and undergraduates with an R background will be better prepared for post-baccalaureate positions, we first and foremost recommend that other ecology instructors use R in their courses when conducting data analysis. We feel that, depending on your course goals, one could take either approach we outlined for our courses and successfully incorporate R into the classroom. Regardless of the approach taken, the following considerations should be made for a successful experience:

1. Be aware that teaching R is different from teaching statistics. In our experience students were weak in statistical skills and, prior to our courses, were unaware of the concepts of reproducibility and documenting steps in data analysis. Students are able to learn R without having a strong background in statistics, but may need some statistical practice in addition to learning the programming language. Sarvary (2014) recommends an approach in which R programming is taught alongside statistics early in a lab section, and then students use both of these skills concurrently throughout the remaining coursework.
2. Be aware of the amount of time it will take to prepare assignments. Preparing and writing assignments was a significant time sink in course preparation. Neither author had previously taught their respective course. As a result, the decision of what to do each week in R plus finding the appropriate datasets and functional packages, or preparing low stakes R assignments, took up more time than preparing lectures for the week. Additional time was spent writing and running the homework and problem set code to ensure it was functional before assigning it to the students. Future iterations of the course will involve less time for assignment preparation.
3. There is a tradeoff between teaching R and teaching content. The instructor should determine the goals and main focus of the course. For example, is data collection and analysis, fieldwork, or theory most important? Determining the priorities of the course is important for understanding how R can be integrated. If fieldwork and data collection are higher priorities, then the approach taken by ELB in her Forest Ecology class would be more appropriate. If the course is taught without a lab, or must cover more theoretical concepts, then LAA’s approach in her Community Ecology class may be a more suitable model. In our experience, it seems that in a typical semester-long ecology course, there is only enough time to do two out of three of the following in enough depth: learning background and theoretical concepts, data collection in the field, or learning statistic principles and/or programming software. For example in CE, the course focused on the first and last of these goals. With an additional lab, perhaps more time could be focused on all three topics, but even in ELB’s case with an extended lab period, time spent in the field and working with R did decrease the amount of time spent on theoretical concepts.
4. Understand the curriculum and how the course fits in. It is very helpful to have a sense of who in one’s institution, and particularly in one’s curriculum, also uses R or is interested in using R. LAA’s and ELB’s courses were complementary to one another in that we each emphasized different uses of R and different aspects of data management. Because of this, students who took both classes could, theoretically, get additional practice, without too much redundancy. An additional, related, recommendation is to determine how much practice students receive with statistics before they enroll in the course being taught, and perhaps even find a way to collaborate with mathematics and/or statistics departments. In our institution, members of the statistics department faculty are willing to work with faculty in other departments on both research and pedagogical projects; this open communication facilitates building students’ skills. Finally, determine if the course must focus on research methods or theoretical background. R can be implemented in both data collection and supporting conceptual material, but may be implemented in different ways (e.g. regular practice with data management similar to ELB’s approach or learn-as-you-go with different packages like LAA’s approach).
5. Identify resources outside of the classroom for students. As most educators know, once students leave the classroom, retention of skills learned diminishes unless they are practiced regularly (Arthur et al 2009). If the goal is to maintain a strong foundation in R programming skills for preparation in post-undergraduate opportunities, then further use after the classroom is necessary. We have helped students further their skills in R programming through senior-year research projects. We have also encouraged students to attend a local R Users Group to learn and practice new skills. Furthermore, advising students to take additional courses that are available, such as an advanced statistics or ecological modelling course, may also be a way to continue their preparation. Both LAA and ELB have found that having taught an R course, we have had students return with R-specific questions about independent research or for seeking advice in coursework to expand their R knowledge. By incorporating R into the curriculum, we have increased the institutional support for learning R and have served as resources for our current and former students.

## Conclusions

In the last decade, R has become the de facto application for data analysis in ecology and its use is increasingly required for post-baccalaureate students joining the ecology workforce or pursuing graduate school. The National Science Foundation “Vision and Change” document (AAAS 2011) identifies several core competencies for undergraduate biology education, three of which can be developed through teaching R: ability to apply the process of science, ability to use quantitative reasoning and ability to use modeling and simulation. As more and larger data sets become available, and as there is a growing push for reproducible research, exposing students to basic data management skills and basic programming in addition to statistics will become even more important. We encourage those instructors who have not yet done so to consider adding some instruction in R to their course designs.

## Acknowledgements

The authors would like to thank the Saint Lawrence University Forest Ecology students from Fall 2017 and Community Ecology students from Spring 2018 for being so open to learning a new and important skill. We’d like to thank participants in the North Country R Users’ Group for discussion and suggestions around using R in the classroom. We’d also like to thank members of the SLU Faculty Writing Group for comments on the manuscript.

