## Appendix S1. for "Teaching R in the undergraduate ecology classroom: approaches, lessons learned, and recommendations"

Appendix S1. R skill development plan and sample R assignments from Community Ecology and Forest Ecology courses.

#### Overview

In the following pages we have put together a week-by-week overview of how we organized teaching R in two undergraduate ecology courses, Community Ecology and Forest Ecology. We have also gathered a variety of sample assignments from each course. First are three example assignments and an example of the final project from Community Ecology, followed by three example assignments and an example of the final project from Forest Ecology.

### Week by week course organization

Table S1. Week by week map of R skill development activities in each course.

|  | R Activities | |
| --- | --- | --- |
| Class Week | Community Ecology | Forest Ecology |
| 1 | Installing and Introduction to R and RStudio | GSWR* preface through chapter 1 |
| 2 | Modeling Competition in R:  1) Learn to plot data using R.  2) Introduce basic R script for interspecific competition models and give an opportunity to test different scenarios | Installing and Introduction to R and RStudio |
| 3 | Calculating Niche overlap using R | GSWR Chapter 2: create .csv files, long vs. wide data formats; import data; install and run packages, introduction to dplyr, ggplot2 packages |
| 4 | Building R Skills with Predator Data:  Tidy, plot, and analyze some fish stomach content analysis data | GSWR Chapter 3 week 1: examine a data set, change column names, basic filtering and grouping of data, basic data summary |
| 5 |  | GSWR Chapter 3 week 2: dplyr and data extraction & summary, factor variables |
| 6 | Building R Skills with Plant Cover Data: Build skills in R to prepare for final project. Examine datasets of plants impacted by volcanic eruptions on Mt. St. Helens (del Moral 2010). There are 3 possible analyses; pick two to complete this assignment.  Analysis 1: Basic mapping  Analysis 2: Regression  Analysis 3: ANOVA and post-hoc analysis | GSWR Chapter 4: basics of ggplot2, box plots, scatter plots, faceted plots |
| 7 | Use Chi-square to examine mutualisms:  Use similar data to Bronstein and Ziv (1997) to conduct a chi-square test to note how a classic mutualism between the yucca plant *Yucca schottii* and its pollinator moth *Tegeticula yuccasella* impacts the presence and activity of two other insect species, the beetle *Carpophilus longus* and the gall moth *Prodoxus y-inversus*. | Problem Set 1 assigned |
| 8 | Plan analysis for and begin final project; meet with Dr. Auker |  |
| 9 |  | GSWR Chapter 5 week 1: introduction to statistical testing; Chi-square and two-sample t test; more ggplot2. Begin formal analysis of class data |
| 10 |  | GSWR Chapter 5 week 2: linear regression, one-way ANOVA |
| 11 | Measuring diversity:  Measure diversity using the Simpson’s Index and the Shannon Index; calculate ‘effective species’ measurement | Continue formal analysis of class data with content learned in GSWR to present |
| 12 |  | Problem set 2 |
| 13 | Dissimilarity index and hierarchical clustering | GSWR Chapter 6: two-way ANOVA |
| 14 |  | Introduction to vegan package; community structure; final project assigned |
| 15 |  | GSWR Chapter 8: plotting tricks and tips with ggplot2 |

*GSWR = Beckerman et al. (2017), Getting started with R: An introduction for biologists.

### Community Ecology Example Assignments

The four Community Ecology assignments include the introductory assignment for learning to use R and RStudio introduced in week 1 (Table S1) as well as the “Modeling Competition” assignment from week 2. The third assignment, from week 6 in the semester, asks students to practice the skills they will need for their final project on a data set related to the eruption of Mt. St. Helens. The fourth example is the final problem set that students were asked to complete at the end of the semester.

#### Community Ecology Example Assignment 1

##### Downloading and Installing R and R Studio

The goal of this assignment is to download and install the software you will use throughout the course and practice uploading a simple dataset so that you are familiar with some very basic commands.

Note: In all assignments, there is room to record errors you have encountered and how you fixed these errors. I recommend that you keep a log of these errors and fixes so that you can recall how to fix them in the future.

##### Part 1: Installing R and R Studio

1) Read pages 1-6 in the *Getting Started with R* (hereafter referred to as “Beckerman”) text. Follow the instructions to download and install both R and RStudio on your computer. (You MUST have R to use RStudio!) RStudio is simply a user interface for R that allows you to see the stored variables, datasets, packages, and other useful things that you cannot easily see while using R itself. From my own experience, it makes learning how to use R much easier!

If you already have R and R Studio installed, make sure they are up to date! To ensure you have the latest versions, do the following:

a) Open up RStudio on your computer.

b) Go to Help, then click on “Check for Updates”. If RStudio is up to date, you will get a dialog box stating that “You’re using the newest version of RStudio.” If not, you should get a dialog box asking if you want to install the latest update. Say yes and follow the instructions for installation.

2) Read pages 6-26 after opening up RStudio. (Note that you never need to open R on your computer as RStudio does this in the background.) Follow the instructions as you read to familiarize yourself with the basics of RStudio. Feel free to play around with it! You can’t break anything, but it is possible you might get errors.

If at any time you get an error, I suggest you record it on the last page of this assignment. Using Google or other sources (see Beckerman section 1.9), try to find a solution to your error (it may be as simple as a missing parenthesis!). Record your solution so you remember how to fix the problem in the future.

##### Part 2: Practice

Now it’s your turn to play with some data! On Sakai, I have posted a simple data set called rpractice1.csv (This file is included in the contents of Assignment 1.)

1) Import your data into RStudio. Here is the simple way to do this (we will get into more advanced methods for inputting R later):

a) Save the rpractice1.csv dataset into a folder somewhere on your computer (don’t forget where it is!)

b) In RStudio, click on “Import Dataset” on the Environment Tab. (Alternatively, you can go to File > Import Dataset…). Choose “From Text (base).”

c) Find the file where you saved it on your computer in step 1 and click “Open”.

d) On the new dialog box, you can see a preview of the dataset. Notice that RStudio automatically adds in generic headers for your columns because it does not know that “SiteNumber”, “Species1”, etc… are headings. To fix this, click on the radio button next to “Yes” in the row designated “Heading.” This will remove the headings V1, V2, etc., and will put the correct headings in the right place.

e) Click “Import.”

f) RStudio will automatically show you your dataset in the upper-left pane. It will also place it in the list of variables on the right hand side of the screen. Notice that RStudio tells you that there are 5 observations of 4 variables. This is important! R automatically assumes that columns are variables and rows are individual cases (or observations, or replicates). We need to remember this for the data sets we will be working with throughout the semester.

2) This file (rpractice1.csv) is a hypothetical dataset showing the abundance of 3 different species at 5 different sites. The numbers under “Species1,” “Species2,” and “Species3” indicate the number of individuals from each species. Let’s get a sum of all of the individuals at each site!

a) Download the rprac.R file from Sakai. The .R extension means that this file is an R Script. You can open it in RStudio by going to File > Open File and finding where you saved this file to your computer.

b) When you open the file, you will see that I have started a script code for you, but you must follow the directions below to finish the script.

c) Basically to find the sum of individuals at each site, we need to add up all of the individuals for each row. This is where things get a little tricky in R. To see this for yourself, you will see the following typed in your script box:

rpractice1[1,2]

rpractice1[2,1]

Follow the instructions in the script to run these lines of code.

**Question 1: What do you think the above lines of code mean?**

Record your answer as a comment as designated in the script box.

Now to find the sum of a row, we need to keep in mind which side of the comma is the row and which one is the column to make sure we’re adding up the right stuff.

Next in the script box, you will see the following:

sum(rpractice1[1,2:4])

**Question 2: What is this code asking R to do?**

In your script, write the code to find the sum of the remaining rows.

**Question 3: How would you modify the code to find the sum of all Species 1 individuals at all sites? (Hint: are you trying to find the sum of a column or a row?)**

When you are finished, save your script (click on the save button). Upload it for credit in Sakai (under “Attachments” when you click on the Assignment). You need to agree to the Honor Code Statement before you can upload your script.

Community Ecology Example Assignment 2
Interspecific Competition

This assignment has four goals:
1) To simulate competitive interactions between r- and K-strategists using a board game, and to learn to plot data collected during gameplay using R.

2) To introduce you to the basics of competition models, beginning with the classic Lotka-Volterra interspecific competition model.

3) To introduce you to basic R script for interspecific competition models and give you an opportunity to test different scenarios.

4) To introduce you to a famous community ecology debate about competition and community patterns.

##### Part 1: Plotting in R

There are actually several ways to plot in R. The text focuses on a more advanced way to plot with a package called ggplot2. Before you tackle this, you will want to read Section 4.2 (pages 80-85) in Beckerman to get a basic idea of the meaning of the code. Once you have done that, download both the kvsr.csv data file and a new R script called competitionplot.r from Assignment 2 on Sakai. Follow the instructions in the script to produce your plot of the data.

You will upload both the completed script and your plot to Sakai for this part of the assignment.

##### Part 2: Competition models

Models, in general, have an important place in ecology as they help ecologists predict the outcomes of species interactions over a longer period of time than can be observed. With the continued collection of empirical data, models are constantly updated and improved to make more accurate predictions.

In order to fully understand ecological interactions, it is imperative that a student in ecology has a basic understanding of some of the classic ecological models and what they mean. Almost every ecology text includes models to explain ecological phenomena. Ecologists, in research presentations and papers, use the language of models. The skill of understanding model equations is transferable to other fields as well, like public health, economics, as examples.

Many students new to models may find it daunting, especially if viewing complex mathematical equations fills one with some anxiety. Not to worry, though, as this assignment will help build your confidence in understanding basic model equations by showing you how to put the equation into words and understanding that each expression is a shorthand way of describing a specific interaction among species.

Let’s start with the Lotka-Volterra interspecific competition model described in your textbook in equations 2.2a and 2.2b (page 29). Below is a general form of these equations written in a way you might find familiar from General Biology:

$$\frac{dN_{1}}{dt}=rN_{1}\frac{(K_{1}-N_{1}-\alpha_{12}N_{2})}{K_{1}}$$

This looks a lot like the logistic growth equation, a model that describes population growth under limited conditions:

$$\frac{dN}{dt}=rN\frac{K-N}{K}$$

Both equations use the following terms, which you should have some familiarity with from General Biology or a previous ecology course:

- *r*: the *per capita* intrinsic rate of increase
- *K*: carrying capacity (how many individuals can be sustained by the environment)
- *N*: number of individuals within the population
- $\frac{dN}{dt}$: change in population size over change in time

The difference between the Lotka-Volterra competition model and the logistic growth model is that the former includes an additional term, α, or the competition coefficient. This coefficient measures how much of one species’ niche/resources is used by the competing species. The more one species utilizes another species' niche, the greater impact it will have on the latter's population growth. This is important to understand. The carrying capacity parameter does not take into account interspecific competition. Therefore, if there is competition, we need to include this additional variable to show that some resources are being shared.

Let’s break down the L-V equation into words to help you understand each component and why it contributes to the change in population of a particular species.

The change in population size over change in time is affected by (1) its *per capita rate of increase* (or, how many offspring each individual can produce), (2) the number of individuals in the population, and (3) how close the population size is to the carrying capacity, including (4) consideration of how much a competing species will use up resources.

Those variables that will negatively impact population growth will be preceded by a minus sign (-), where naturally those variables that increase growth are positive. Both *r* and α affect population growth in a way that is dependent on the number of each species within the community.

##### Part 3: Simulating r- and K-strategist competition in R

I have written a script for simulating competition between r- and K-strategists using the Lotka-Volterra model. Download the file lvcompmodel.R and open it in RStudio. Carefully look it over, being sure to read the descriptive comments, explaining each line of code. You may not understand every piece of each line of code, and that’s okay. But it should give you some idea of what you need to do to run a model in R (or in any other type of simulation software). This code:

- Provides the model equation (as done within the created function lvcomp2).
- Gives parameters for the simulation (in this case, it is setting values for *r, K,* and α for each species.)
- Gives starting conditions (as mentioned in the comments, initial population sizes at the start of the simulation).
- Sets length of time for the model to run (done within the parameters of the **ode** function).

Board game simulation in R

Now, let’s run a simulation similar to that run on the board game. Change the parameters (“parms”) to the following:

r1 = 1, r2 = 0.4, k1 = 37, a12=0.7, k2 = 37, a21=0.3

Let’s figure out where these values come from. Since *r* is per capita increase, this is the number of offspring each tree can put out. For the r-strategist in our game, every tree had the potential to produce one seed per turn in our game. The K-strategist, however, could only produce a seed in medium and large trees, which are 40% of the available trees. The carrying capacity is estimated here by the number of spaces on the board. I assigned the alpha values by estimating sunlight absorbed by each tree species (perhaps there is a better way to estimate alpha?). By the end of the game the K-strategist is getting 70% of the sun's energy and the r-strategist gets about 30% of sun's energy.

Make sure the “times” value is set to 1:18 in the ode function, and that the initial conditions ("initialN") are set to c(2,2).

Run the model (you can simply highlight the whole script and click “Run”) and then answer the questions that follow.

**Question 1:** Which species (1 or 2) is the r-strategist and which species is the K-strategist? How did you determine this from the parameters you set? Does the graph plotted from the model match the graph you plotted in Part 2?

**Question 2:** Change the ‘times’ parameter to 1:100 (or 100 time units). What eventually happens to species 1 and species 2? Why?

Let’s run some more simulations. In the chart below, there are new parameters to enter into the model. All other factors (initial starting conditions and time) will stay the same (“initial” = c(2,2) and “times” = 1:100). Run the simulation from the **lvcompmodel** script after changing the parameters in each scenario, then note which species wins. Record the result by checking the appropriate box.

| Scenario | r_1_ | r_2_ | K_1_ | K_2_ | α_12_ | α_21_ | Species 1 wins | Species 2 wins | Both species coexist |
| --- | --- | --- | --- | --- | --- | --- | --- | --- | --- |
| 1 | 0.5 | 0.5 | 1300 | 1500 | 1 | 1 |  |  |  |
| 2 | 0.5 | 0.5 | 1500 | 1500 | 1 | 0.7 |  |  |  |
| 3 | 0.5 | 0.5 | 2100 | 1800 | 0.5 | 0.9 |  |  |  |
| 4 | 0.5 | 0.5 | 700 | 850 | 0.3 | 0.4 |  |  |  |

**Question 3:** For each scenario, explain briefly why the outcome of the simulation occurred from an ecological perspective, based on the parameters given above.

##### Part 4: Reading about competition by two classic community ecologists – a debate!

For Tuesday, January 30, read the assigned paper based on your team number. The person who was the K-strategist player will be the discussion leader for their paper.

Team #1: Diamond and Gilpin (1982)

Team #2: Connor and Simberloff (1979)

For Tuesday, be prepared to answer the following questions in discussion:

1. Where does your paper stand on the chance vs. competition debate?

2. What problems did your paper’s authors have with previous researchers’ claims?

3. Are the arguments within the paper that you read convincing? Why or why not?

Responsibilities of the discussion leader:

1. Ask one additional question specific to your paper other than those listed above. Record your question and a summary of the group’s answers on a piece of notebook paper and hand in at the end of class.

2. Summarize the paper to the rest of the class.

Error Log

| Error | Solution |
| --- | --- |
| Example:  Error: unexpected ‘)’ in: “sum(rpractice1[1, 2:4)” | Example:  Forgot closing bracket. Insert before closing parenthesis. |
| Example:  Error: object ‘rpractice’ not found | Example:  Mistyped the name of the dataset. Add in ‘1’ after ‘rpractice’. |

#### Community Ecology Example Assignment 3

For this assignment, we are going to take a look at datasets of plants impacted by volcanic eruptions on Mt. St. Helens. These datasets came from del Moral (2010). The purpose of this assignment is to build your skills in R to prepare you for your final project. There are 3 possible analyses that you can do on this dataset. Pick TWO of these analyses to complete this assignment. Your best bet is to pick analyses that are most relevant to your final project or that are most interesting to you.

Here are the analyses and the page numbers within this document where you can find them.

Analysis #1: Basic mapping skills	Page 1

Analysis #2: Regression	Page 4

Analysis #3: ANOVA and post-hoc analysis	Page 5

Each analysis section will describe what you need to submit for credit for this assignment.

##### Analysis #1: Basic mapping skills

Sometimes, you just want a simple map for a publication or presentation that doesn’t have the “bells and whistles” of a GIS map. (However, GIS skills are useful for an ecologist and I recommend you get a basic foundation in those skills as you continue to pursue fields in ecology and conservation biology.) The good news is that R can produce maps, even colorful maps based on readily available platforms like Google, to show your data. (Try doing that in Excel!)

The purpose of this part of the assignment is to create a simple map of plot locations within the Mount St. Helens sampling area. I am going to show you how to do this, using high-resolution spatial polygon data available from the web.

Follow the following steps to create your map:

1. Install and load the following packages in RStudio. (Use what you’ve learned in previous assignments to do this!): raster, ggmap.
2. We are going to create two different maps: a simple line map of the county in which Mt. St. Helens is located and another of the volcanic park itself, using a Google Map. We will plot points on both maps. Sometimes a simple line map may be clear enough to show political boundaries, but other times you may want something “prettier” to show off certain details, like terrain, roads, or other features.

For the line map, we are going to use the ‘raster’ package to obtain spatial polygon data available on the web and focus in on the county in which the volcano is located.

So, using the Internet, in what state and in what county can you find Mount St. Helens?

3. In a new script in R, use the following line of code to retrieve and save spatial data for the United States.

us <- getData(‘GADM’, country=’USA’, level=2)

Note: The level refers to how much detail your map has. You can see this yourself if you

enter and run the following line of code:

plot(us)

You can see that this gives you a map of every county in the US.

At the moment this map is very small and difficult to see. We need to zoom in on our location of interest.

4. Run the following code to see what data are actually available in our ‘us’ data frame.

names(us)

The resulting output gives you various names in the spatial polygon data frame we have named ‘us.’ Specifically, you may be interested in NAME_1 and NAME_2. NAME_1 represents state names, and NAME_2 represents county names. You can see this by running the following lines of code:

unique(us$NAME_1)

unique(us$NAME_2)

Let’s make two maps: one of the state in which Mt. St. Helens is in and another with the county in which the volcano is located. I will explain how to make a map of the state, but will challenge you to create and run the code to make a county map on which we will plot points!

5. Make a map of the state.

Using the following code, and inserting the correct state where indicated, plot a map of the state where you can find Mt. St. Helens.

state<-subset(us, NAME_1==”InsertStateHere”)

plot(state)

6. Make a map of the county.

Change the code above to plot the *county* where Mt. St. Helens is found.

7. Plot points on your county map.

Now, let’s plot some points from our dataset onto this county map. Download and import the “mshplot.csv” dataset into RStudio.

In this dataset, there are two columns you should be interested in right now: the LONG and LAT columns. These give the longitude and latitude of each of the sampling sites. Let’s plot them on your county map. Make sure that the county map is the last thing you have in your Plots tab. If you played around with the code and plotted something else, please rerun the code from step 6 before continuing on.

Simply run the following code:

points(mshplot$LONG, mshplot$LAT)

Congratulations! You have plotted your first set of points on a map in R! Export and Save this plot as an Image. Call it: Analysis1LineMap. You will submit it as part of this assignment.

8. As you can see, while you have plotted points on the map, this map is pretty uninformative: no legends, no labels, kind of boring, right? The good news is you can use maps from Google right in R using the ‘ggmap’ package. Now we can see some useful details. And, I will also teach you how to vary the points on the graph so they are more informative.

Run the following codes to get a map of our campus:

You can basically make a map out of anything using this technique. Try making a map of SLU:

slu<-get_map(location=”St. Lawrence University”, zoom=15)

slumap<-ggmap(slu)

slumap

Cool, huh?

Now, make a map of Mount St. Helens by changing the code above. This is a rare case where capitalization and format do not matter in R since you are basically searching for the location on Google.

9. Plotting points on your Google Map of Mount St. Helens.

Now comes the interesting part. The package ‘ggmap’ is based on ‘ggplot’ so the syntax is similar to what is found in your Beckerman text. Let’s try plotting our points on the Mt. St. Helens map. Note that in my code below, I called my map ‘mshmap’. If you called your map something different in step 8, you should change this in the code below:

mshmap+geom_point(aes(x=LONG, y=LAT), data=mshplot)

Now you have a plot of your sites! But we are not done yet! Notice that in the mshplot data, the sites differ by succession type. What we are going to do now is change the colors of the points based on the succession type. Run the code below (note the difference from the code above).

mshmap+geom_point(aes(x=LONG, y=LAT, colour = SUCCESSION_TYPE), data=mshplot)

Now you have a useful map of your sites distinguished by the type of succession shaping the community at each site.

Export and Save this plot as an Image. Call it: Analysis1SuccessionMap. You will submit it as part of this assignment.

For this analysis, you should submit these two files to Sakai:

1) Analysis1LineMap.jpeg

2) Analysis2SuccessionMap.jpeg

NOTE: For those of you who want to make maps for your project, I recommend the ‘Geospatial Analysis in R’ link listed under References at the end of this document. Here, you can learn to make heat maps, too!

##### Analysis #2: Regression

You should do a regression if you want to see if two continuous variables correlate with one another.

With our data, we are going to ask the question: Does elevation affect plant percent cover on Mt. St. Helens?

Follow the steps below to answer this question:

1. Read pages 109-118 in the Beckerman *Getting Started with R* book.
2. Download and import the “plantcover.csv” dataset into RStudio.
3. Create a graph similar to figure 5.6 in your book (with line and standard error plotted on the graph) with Plant Percent Cover on the y-axis and Elevation on the x-axis. Export and Save this Graph as Analysis2.jpeg. You will submit this in Sakai.
4. Copy and paste the table below into a new Word Document and save it as Analysis2.doc. Fill in the table with the results from the regression analysis (using the anova and summary functions). Submit this document in Sakai.

| F Value (from anova() function results) |
| --- |
| Pr (>F) (from anova() function results) |
| Estimated intercept |
| Estimated slope |
| Does elevation affect plant percent cover on Mt. St. Helens based on your findings? |

For this analysis, you should submit these two files to Sakai:

1) Analysis2.jpeg

2) Analysis2.doc

##### Analysis #3: ANOVA and post-hoc analysis

You should do an ANOVA if you are comparing the means of more than 2 samples.

With our data, we are going to ask the question: Does mode of succession impact plant percent cover on Mt. St. Helens?

Follow the steps below to answer this question:

1. Read pages 118-127 in the Beckerman *Getting Started with R* book.
2. Download and import the “mshanova.csv” dataset into RStudio.
3. Create a graph similar to figure 5.8 in your book (use coordflip()!) with succession on the y-axis and total cover on the x-axis. Export and Save this Graph as Analysis3.jpeg. You will submit this in Sakai.
4. Copy and paste the table below into a new Word Document and save it as Analysis3.doc. Fill in the table with the results from the ANOVA analysis (using the anova and group_by()/summarise() functions). Submit this document in Sakai.

| F Value (from anova() function results) |
| --- |
| Pr (>F) (from anova() function results) |
| Is there a significant difference in plant percent cover in different modes of succession on Mount St. Helens? |
| What are the mean percent cover values for each succession mode? |

For this analysis, you should submit these two files to Sakai:

1) Analysis3.jpeg

2) Analysis3.doc

#### Community Ecology Project Assignment

Goal: To analyze, graph, and describe the significance of a community ecology dataset.

Due: Final poster file due by 5pm on April 22, 2018; Presentation day: In class 4/26

There are two parts of this assignment to reflect the two-part learning goals of Biol 4021:

1) To analyze and plot/map a dataset using R/RStudio
2) To qualitatively describe the system and interactions within the given community described in your dataset.

The poster will contain:

- Introduction:
  - Description of taxonomic groups within data set
  - Description of study area and biome type
  - Questions you came up with about the dataset
- Methods
  - How the data were collected
  - Sample R script used to conduct graphical and statistical analysis
- Results
  - 1+ graphs showing relationships within dataset
  - Statistical results
- Discussion of community interactions, patterns, and processes; what future experiments would help clarify potential relationships or answer questions that resulted from this analysis?
- Literature cited, including paper sourced for data

Milestone Dates for Project

Week of Jan 30 – Topics/biomes assigned

Week of Feb 20 – Meet with Dr. Auker to discuss questions, approaches

Week of March 27 – Share graphical analysis and results

April 22 – Poster must be completed and sent to printer by this date

April 26 – Presentation of posters in class

What to do next:

When you receive your dataset, examine the data and come up with ways to analyze it. Reading the paper will help you formulate your questions. These questions must meet the following criteria.

- One analysis must focus on statistical relationships (e.g. ANOVA, t-test, regression, etc.)
- One analysis must focus on diversity and/or community patterns
- The final analysis must be graphical: either a map or a plot.

To help you start brainstorming, answer these questions: What analyses make sense based on your data? What interactions may occur within this community? How can you test for these relationships?

You must complete your poster and submit it via this link by 5pm on April 22 (NO EXCEPTIONS due to the quantity of requests going through the LSL GIS lab at this time!!!): <http://www.stlawu.edu/library/large-format-printing-services-launders-science-library>

Your poster size will be 24 x 36 (Yes, it’s not a huge poster, but should be enough to showcase your findings).

Follow the following guidelines for creating your poster: <http://www.stlawu.edu/library/content/poster-design-guidelines>.

We will have an in-class symposium (with some invited guests, perhaps) on April 26^th^ to check out what you have done. You will also have an opportunity to check out each other’s work on that day. More details TBA.

### Forest Ecology Example Assignments

The four Forest Ecology assignments include three examples of "low stakes" assignments that were assigned alongside Chapter 3 from Getting Started With R: An introduction for biologists (Beckerman et al. 2017) during weeks 4 and 5 of the semester (Table S1). The fourth example is the final problem set that students were asked to complete at the end of the semester.

#### Forest Ecology Example 1

GSWR Chapter 3 Exercise 1

Work through the material in GSWR Chapter 3, not including Appendix 3A or 3B. Create a R script that contains all of the material presented in the chapter. Be sure to read through Table 3.1 on p. 64. Add the following features to your R script.

1. Save your script as LastnameFirstInitial.Ch3Ex3.R
2. Add the following commented lines at the top of your script:

#FirstName LastName

#Chapter 3 Exercise 3

3. The rest of the code should be whatever you need to work with the compensation dataset from the text and the steps that the chapter follows.

Submit your R script to me via email.

By the end of this exercise, you should be comfortable with the following functions and what they do:

- select()
- slice()
- filter()
- arrange()
- mutate()
- with()
- group_by
- summarise()
- mean()
- sd()

Send your R script to me via email.

#### Forest Ecology Example Assignment 2

GSWR Chapter 3 Exercise 2

This exercise is designed to help you reinforce concepts from GSWR Chapter 2 and Chapter 3 and to push you to apply functions from the dplyr package more independently.

Please do the following:

1. Open R Studio server (rstudio.stlawu.local:8787)
2. Start a new R script file and save it as “Ch3Ex1.R”. Comment the beginning of the R script with #FirstName LastName and #Chapter 3 Exercise 1
3. Add code to turn on the dplyr package and code to “clear R’s brain.”
4. Import the csv file called “donnerville_percent_cover.csv” located in rstudioshared/Biol4018_F17_ForestEcology/GSWR/Data.
   a. Name the dataset “Cover” during import
   b. Be sure to copy the first 2 lines of the import code and paste them in your R script.
5. Examine the data set using at least two different functions.
6. Consider renaming column headings; this choice is up to you.
7. Determine the average percent cover by cover type (bare, bryophytes, vascular plants) for Donnerville 1.
8. Determine the average percent vascular plant cover across all sites.

Send your R script to me via email.

#### Forest Ecology Example Assignment 3

GSWR Chapter 3 Exercise 3

This exercise is designed to help you reinforce concepts from GSWR Chapter 2 and Chapter 3 and to push you to apply functions from the dplyr package more independently.

Please do the following:

1. Open R Studio server (rstudio.stlawu.local:8787)
2. Start a new R script file and save it as “Ch3Ex2.R”. Comment the beginning of the R script with #FirstName LastName and #Chapter 3 Exercise 2
3. Add code to turn on the dplyr and ggplot2 packages and code to “clear R’s brain.”
4. Import the csv file called “donnerville_tree_tally.csv” located in rstudioshared/Biol4018_F17_ForestEcology/Class exercises.
   a. Name the dataset “Tally” during import
   b. Be sure to copy the first 2 lines of the import code and paste them in your R script.
5. Examine the data set using at least two different functions.
6. Consider renaming column headings; this choice is up to you.
7. Create a new dataset called “Trees” that includes all of the data for all of the sugar maple, white ash, basswood and American beech observations.
8. Now determine the count for each of these 4 species in each of the height classes. (Hint: This one is difficult: remember that you can call more than one variable in a group_by function and also consider using arrange()).
9. Plot the number of trees in each height class in each species. Hint: look at?qplot and see how they use the color call in the examples).

Send your R script to me via email.

#### Forest Ecology Final Problem Set

FOREST ANALYSIS FINAL ASSIGNMENT

At this point in the semester, we have finished gathering all of our forest data and have worked through all but 2 chapters (7, 9) in GSWR. In problem set 2, you began your forest analysis by examining how our four forests differed in terms of ground cover and litter depth, and in terms of the number of seedlings and trees (as well as their density and species composition) among forests. All of those steps were elements of your analysis; you may wish to include some of them here.

For this assignment, I’d like you to accomplish two broad goals:

1. Get experience doing your own data analysis with little guidance from me (but hopefully based heavily on the examples we’ve done in class)
2. Take time to visually examine and think about the biology of our forests.

To that end, for the final forest analysis assignment, please do each of the following steps:

1. Choose *one forest plot* from among the 12 we measured (not one forest, but a single plot from one forest) and use your R skills (both in terms of summarizing data and in visualizing data) to DESCRIBE the plot. I’m interested that you describe the plot in terms of the data, and NOT in terms of e.g. the plot descriptions we wrote out at the start of the semester.

Your description should include:

a. The R code (in your R script or scripts)
b. An overview narrative of a few sentences to a paragraph in your word document orienting me to the first section
c. Organized and properly labeled table or tables, as you choose, embedded in a word document
d. Neatly organized and properly labeled figure or figures, as you choose, embedded in a word document
e. Summary statements ABOUT the data (not describing the data, but describing any trends that you think are interesting or worth pointing out) in the word document.

2. Choose *one forest* (including 3 plots) from among the 4 forests we measured and use your R skills (both in terms of summarizing data and in visualizing data) to describe and possibly analyze the Forest.

Your description should include:

a. The R code (in your R script or scripts)
b. An overview narrative of a few sentences to a paragraph in your word document orienting me to the second section
c. Organized and properly labeled table or tables, as you choose, embedded in a word document
d. Neatly organized and properly labeled figure or figures, as you choose, embedded in a word document
e. Summary statements ABOUT the data (not describing the data, but describing any trends that you think are interesting or worth pointing out) in the word document.
f. For this level of analysis, you may also wish to run a statistical test or two to examine differences you may observe among plots within your forest. Doing so is not required, but is allowed if it helps you tell an interesting story with the data.

3. At the level of the *comparing among forests*, please provide the following:
   a. R code (in terms of R script or scripts) for any data manipulations or analyses;
   b. An overview narrative of a few sentences to a paragraph in your word document orienting me to the third section
   c. A figure comparing the Shannon diversity index among the four forests, embedded in the word document;
   d. Results of a one-way ANOVA, including Tukey HSD comparisons if necessary, and summarized in the word document, testing if there is a significant difference in the mean Shannon index among the 4 forests. Please use the Shannon index that is based on both tree and seedling diversity.
   e. A correlation test that correlates the many environmental variables we have collected among our forests and a plot of the correlation tests showing the magnitude and significance of the correlations, included in the word document
   f. A faceted plot illustrating, for each forest, the importance value of each species (so facet by forest)
   g. An ordination plot based on overall importance values that aligns the tree species and our forest plots along two ordination axes
   h. A second ordination plot that shows the strength and direction of any environmental factors that are significant at P<0.05 or lower.
   i. If any environmental variables are identified in the ordination process as contributing to separation of the forests, include for up to but not more than two of the environmental variables a boxplot showing their values compared among forests and output from a one-way ANOVA testing to see if the mean of the variable differs among forests.
   j. Make sure that each plot or table you include in your analysis is well-labeled, nicely designed, and is followed by a descriptive sentence or two highlighting what you think is important to see.
   k. A landscape-level comparison of your choosing, along with associated tables or figures.

Please keep in mind as you work that I will grade your assignment based upon:

1. The degree to which you follow these instructions
2. The degree to which your R code produces mathematically correct answers
3. The degree to which you clearly annotate your R code so that I can tell what you were thinking/doing when I work through it
4. The care you take in putting together your word document (did you include table headings, figure legends, clearly written/thought out reflections on the trends in the data)
5. The degree to which your code and steps follow the basic workflow we outlined in Problem Set 2 and that appears again below
6. The care you take with making nice-looking plots that are as effective as possible in conveying your message (did you spruce them up using tips from Chapter 8 to improve their effectiveness?)

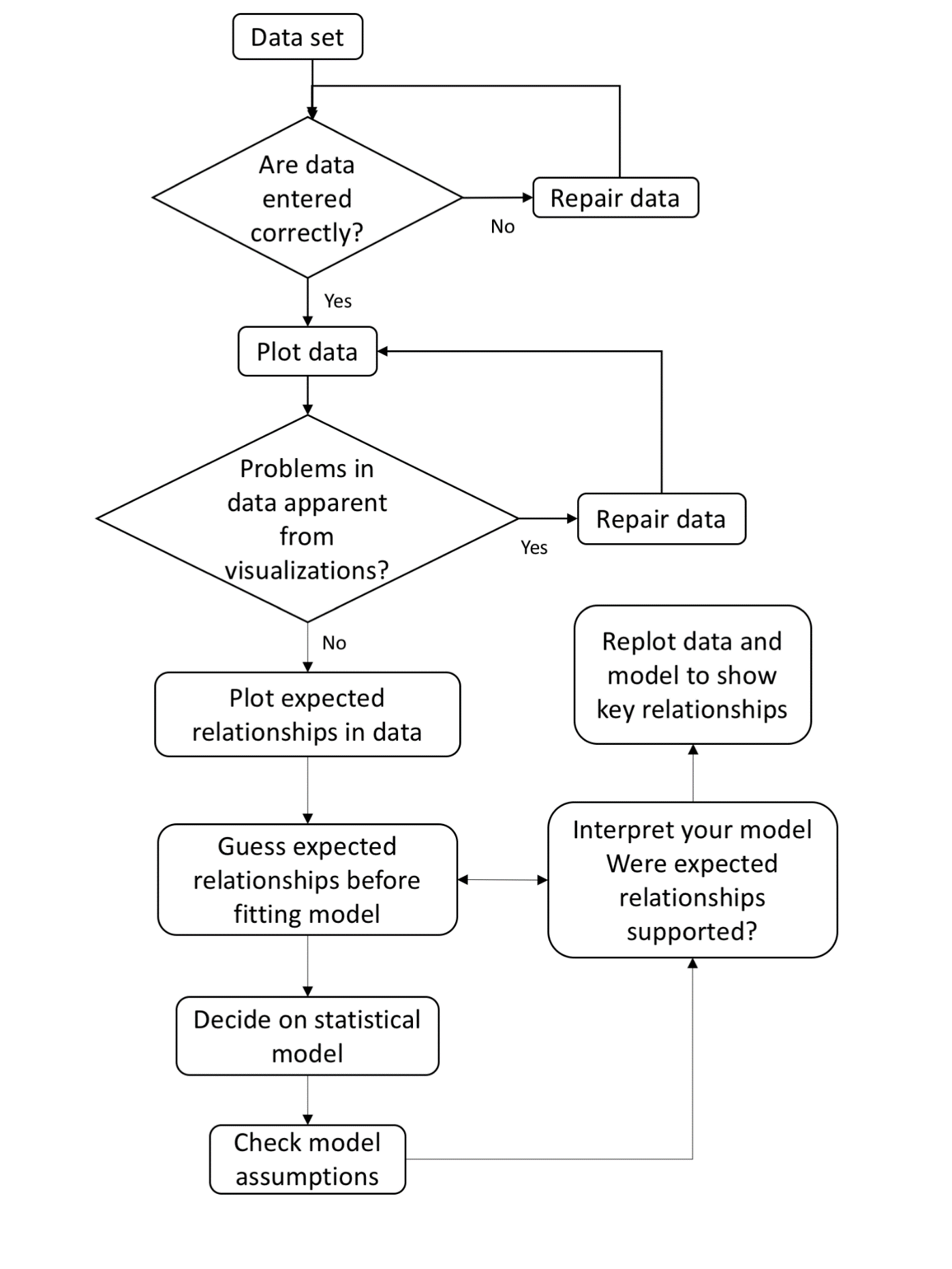

Guidelines:

A. The data you will need are located on the T drive in a folder at t://Barthelmess/FE2017/Forest Analysis. There are multiple csv and excel files there with data for you to choose from.
B. I have not intentionally added any error to the data.
C. Please write your RScript using RStudio server, or at least make sure that when you load the data in your R script, it calls a dataset located on the Rstudio server.
D. Make sure any .R scripts you submit have all of the “default” code that we have been including in every .R script this semester.
E. Note that our forest plots are 400m^2^ which is = 1/25 ha. Use a scaling factor of 25 to get density of trees. Our regeneration plots are 0.001 ha – use a scaling factor of 1000 to get density of seedlings.
F. You will submit a word document that answers the questions, one or more .R scripts (your choice) as well as any images you produce, all collected together in a .zip file. If you are uncertain how to zip files together, see the page on our Sakai site that explains it. You will then email me the zipped file with all your work. Name the zipped folder “LastnameForestAnalysis.zip”

mutualism. *Oecologia* 112(3): 379-385.

del Moral, R. 2010. Thirty years of permanent vegetation plots, Mount St. Helens, Washington.

*Ecology* 91: 2185.
